# Pixel-based machine learning and image reconstitution for dot-ELISA pathogen serodiagnosis

**DOI:** 10.1101/2020.03.18.997320

**Authors:** Cleo Anastassopoulou, Athanasios Tsakris, George P. Patrinos, Yiannis Manoussopoulos

## Abstract

Serological methods serve as a direct or indirect means of pathogen infection diagnosis in plant and animal species, including humans. Dot-ELISA (DE) is an inexpensive and sensitive, solid-state version of the microplate enzyme-linked immunosorbent assay, with a broad range of applications in epidemiology. Yet, its applicability is limited by uncertainties in the qualitative output of the assay due to overlapping dot colorations of positive and negative samples, stemming mainly from the inherent color discrimination thresholds of the human eye. Here, we report a novel approach for unambiguous DE output evaluation by applying machine learning-based pattern recognition of image pixels of the blot using an impartial predictive model rather than human judgment. Supervised machine learning was used to train a classifier algorithm through a built multivariate logistic regression model based on the RGB (“Red”, “Green”, “Blue”) pixel attributes of a scanned DE output of samples of known infection status to a model pathogen (*Lettuce big-vein associated virus*). Based on the trained and cross-validated algorithm, pixel probabilities of unknown samples could be predicted in scanned DE output images which would then be reconstituted by pixels having probabilities above a cutoff that may be selected at will to yield desirable false positive and false negative rates depending on the question at hand, thus allowing for proper dot classification of positive and negative samples and, hence, accurate diagnosis. Potential improvements and diagnostic applications of the proposed versatile method that translates unique pathogen antigens to the universal basic color language are discussed.

## Introduction

Reliable and timely diagnosis of disease-causing pathogens is of paramount importance for maintaining optimal health for people, animals and the environment under the integrated One Health approach (***World Health Organization, 2018***). One of the most widely used assays for pathogen detection is the enzyme-linked immunosorbent assay (ELISA); in its solid-state version, dot-ELISA (DE), samples are directly applied to or dotted on a nitrocellulose or nylon membrane and then probed for detection by specific antibodies in a chromogenic enzymatic reaction. DE is a rapid screening test that is at least as sensitive as ELISA, yet it is much cheaper and requires no special equipment or working conditions. As a result, DE has been adopted in a wide array of applications, ranging from disease diagnosis in humans (***Rodkvamtook et al., 2015; Subramanian et al., 2016***), animals (***Fisa et al., 1997***) and plants (***Savary et al., 2012***) to microbe and toxin detection in foods (***Venkataramana et al., 2015***).

DE is a qualitative method designed to yield either a positive or negative result in a binary mode. The assay follows the ELISA architecture with similar direct or indirect versions that employ primary and secondary, enzyme-conjugated monoclonal or polyclonal antibodies for targeted antigen detection. In most DE applications, dots are colored by a formazan/indigo dye (***Smejkal and Kaul, 2001***), an *in situ* complex precipitate produced by alkaline-phosphatase (AP) 5-bromo-4-chloro-3-indolyl phosphate/nitro blue tetrazolium (BCIP/NBT) substrate degradation. Output evaluation after color development is empirically eye-based; consequently, positive sample recognition is subjective (***Pappas, 1994; Waner et al., 1996***) and particularly error-prone. Such errors may stem from overlapping dot colorations resulting from host pigment contamination (***Chen et al., 2012; Olkkonen and Ekroll, 2016***), or from the inherent color discrimination thresholds concerning saturation and hue of the human eye (***Baraas and Zele, 2016; Krudy and Ladunga, 2001; Olkkonen and Ekroll, 2016; Reinhard, 2008***).

Two types of errors are possible in binary classification tests: false positives (FP) or “false alarms” and false negatives (FN) or “missing values,” referring to the incorrect recognition of true negatives (TN) as positives and true positives (TP) as negatives, respectively. Proper classification of TP and TN as such is known as sensitivity and specificity, correspondingly (***Trevethan, 2017***). The two error types are inversely related, and their rates must be decided on a case-by-case basis since the arbitrary choice of one over the other could have significant epidemiologic and/or economic repercussions. For example, the detection of quarantine or dangerous pathogens by diagnostic screening tests may require the adoption of a low cutoff and high sensitivity, which could yield increased numbers of FP, but it would also increase the probability of identifying all pathogens or infected individuals (or reduce the probability of missing any), thereby preventing the release of the pathogen to the environment or to uninfected Individuals or populations. In contrast, selectively detecting a common pathogen in the presence of related species or unrelated signals (background noise) may necessitate the adoption of a high cutoff and specificity, thus reducing FP rates and costs associated with additional unnecessary diagnostic tests.

Analogously, accepting *a prior*i as positives only the evident dark spots in a DE readout will result in a high rate of missing values (FN, leaving TP undetected), whereas accepting most or all colored dots as positive will result in high rates of FP (negatives mistakenly considered positives). Either erroneous classification will influence the outcome of diagnosis. A reliable method is therefore needed for non-subjectively discriminating positive and negative samples in DE outputs. Until now, apart from some procedural attempts for improving dot quality either by removing unspecific pigmentation in plant samples (***Chen et al., 2012***), or by controlling the enzymatic reaction time during incubation (***Lathwal and Sikes, 2016***), no work has been done on objectifying and improving DE output interpretation. Artificial intelligence has not been explored for DE output evaluation or FP/FN cutoff estimation. Herein, we describe an image pixel-based supervised machine learning method for DE output evaluation. To the best of our knowledge, applying machine learning to accomplish this goal using an impartial predictive model rather than human judgment is novel.

## Results

### The new approach for DE output evaluation: from pixel information to dot classification and diagnosis

Our approach to evaluating DE outputs and predicting the infection status of samples followed five steps (***Figure 1***). First, a dot blot of a prototypic DE dataset that included known positive and negative control samples was scanned at a resolution of 150 dpi and a “Pixel Information Matrix” of this prototypic image was constructed with the X-Y position coordinates and RGB color information for the pixels of known infection status dots. This process was followed by supervised training of a classifier algorithm to predict pixel probabilities based on a random subset of the prototypic image pixel information. The third step included the cross-validation of the trained classifier algorithm by predicting pixel probabilities in a random subset of the original dataset not used for training, while the fourth involved confusion matrix and Receiver Operating Characteristic (ROC) curve analysis and selection of the appropriate cutoff value. Using the trained and validated classifier, pixel probability predictions could be made after matrix construction of DE readouts of unknown samples. DE image reconstitution of unknown samples at the selected cutoff allowed for proper dot classification and, hence, accurate diagnosis. The entire process, after running the blot using standard methods, could be completed in a few seconds.

**Figure 1.**
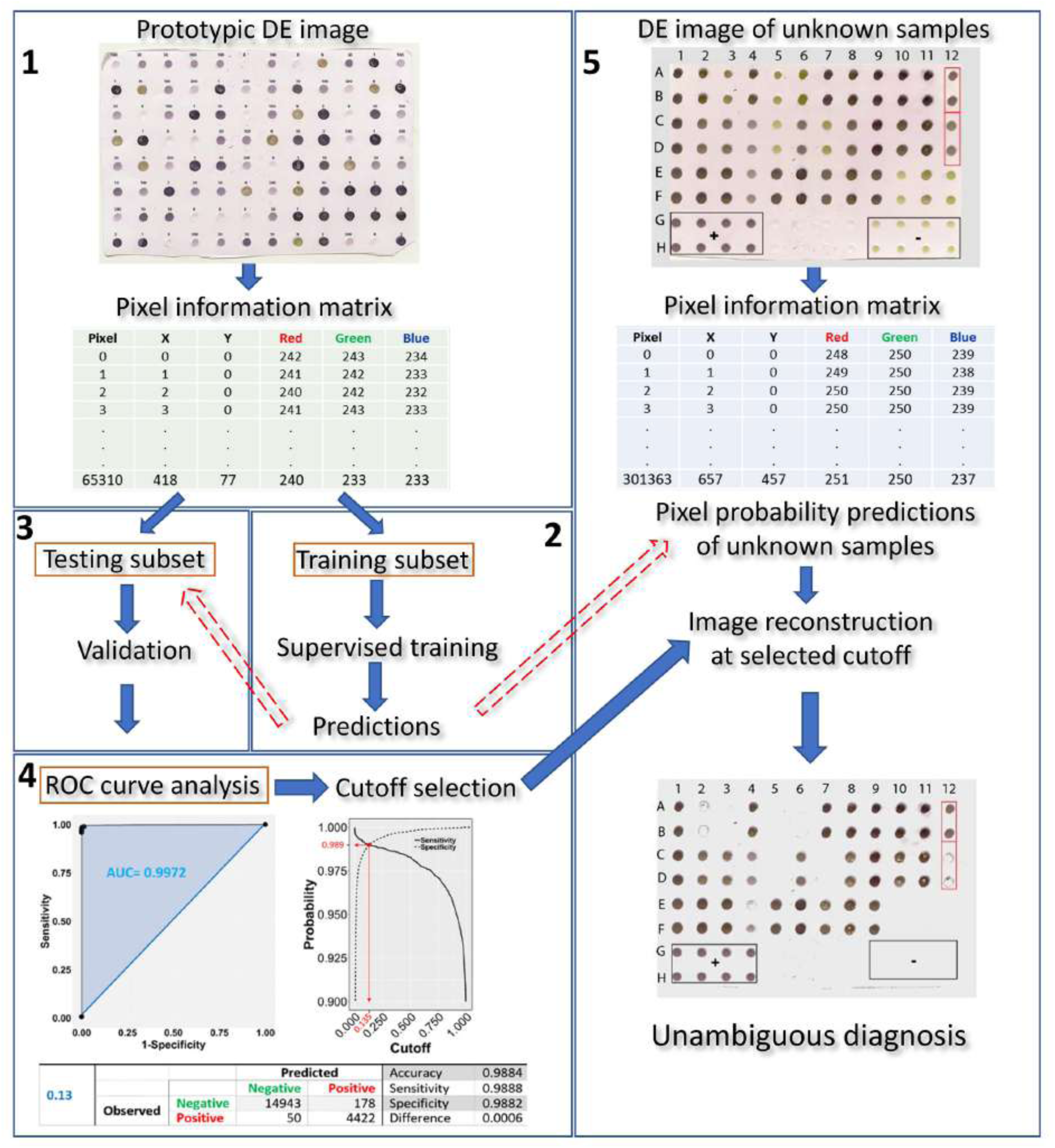
Overview of the proposed machine learning approach for DE output evaluation that harnesses pixel information for dot classification and diagnosis in merely a few hundreds of seconds after running the blot using standard methods. The DE output is first scanned (at a resolution of 150 dpi) and the prototypic image is converted to a matrix, holding pixel position and color information for all pixels contained in the dots (1). The matrix is then randomly tessellated into a training and testing dataset at a 70:30 ratio. Next, training of the classifier algorithm (2) is supervised by a logistic model of choice which, following validation (3), is used for infection status predictions on the testing subset. ROC curve analysis allows for selection of an appropriate cutoff (4). The procedure of scanning the blot and harnessing its pixel information is repeated on a DE of unknown samples (5). Pixel probabilities of the unknown samples are predicted by the trained classifier and a new image of the blot, with dots consisting of pixels with probabilities above the selected cutoff, is reconstituted, allowing for proper dot classification and, hence, unambiguous diagnosis.

### Harnessing pixel information of the prototypic image to train the classifier algorithm

Negative controls in the prototypic DE output displayed a light olive-green color, readily distinguishable from the colors of other categories of samples by the naked eye (***Figure 2***). However, the color of positive controls ranged from dark purple in undiluted or low sample dilutions to light violet at higher sample dilutions, with lighter colorations at higher dilutions. Color hues tended to overlap between similar level dilutions, either at the high or low end (i.e., 1:50 *vs*. 1:100 or undiluted *vs*. 1:2, respectively), rendering eye-based dot discrimination uncertain.

**Figure 2.**
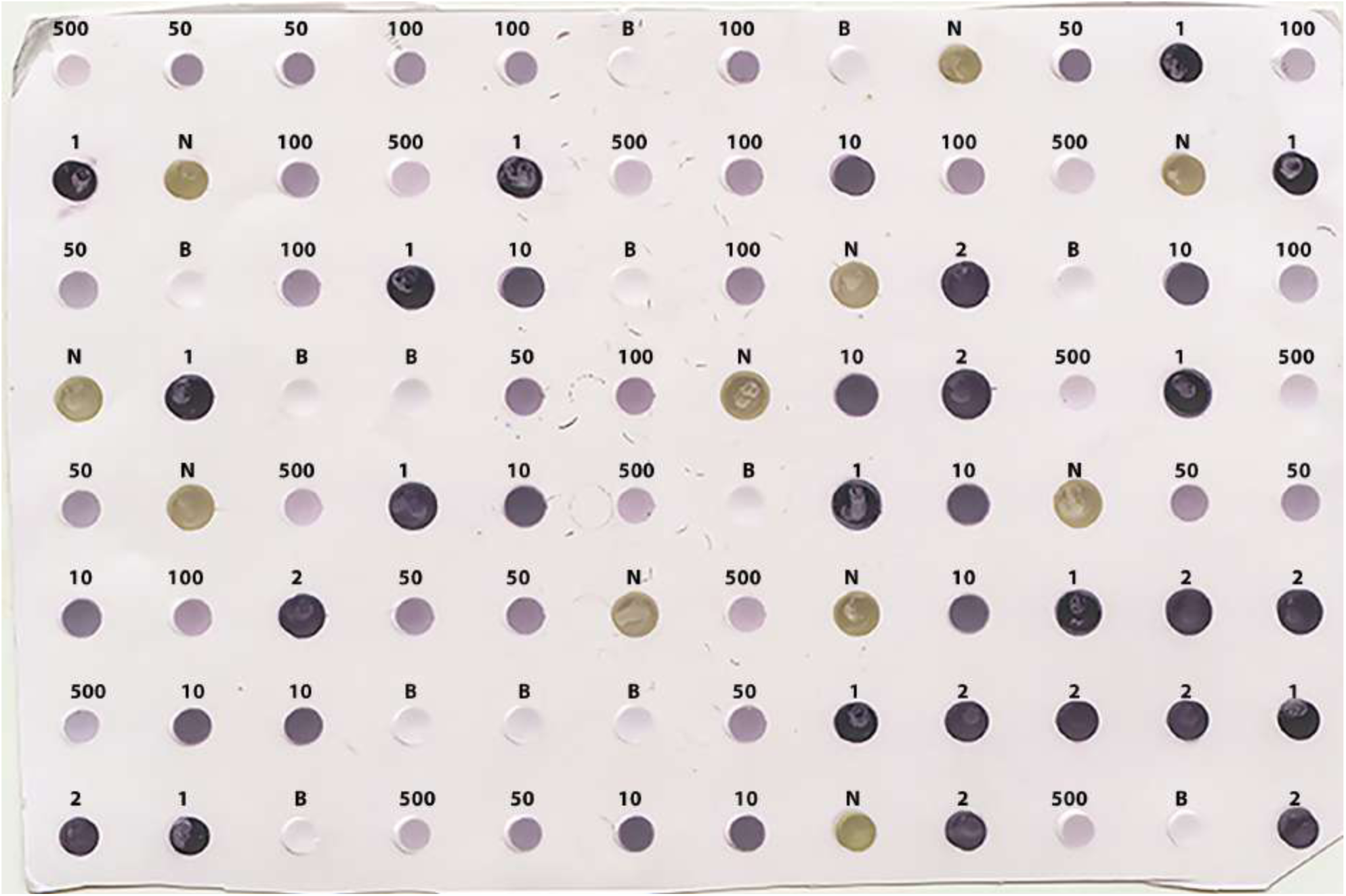
Scanned image (at 150 dpi resolution) of the prototypic DE output used for training the classifier algorithm. Positive, negative and buffer controls were randomly distributed on the membrane in 12, 11 and 13 replicates, respectively. The letters on top of the dots denote the Buffer (B) and the negative (N) controls, while the numbers 1, 2, 10, 50, 100, and 500 denote no dilution, or 1:2, 1:10, 1:50, 1:100, and 1:500 dilutions of the positive control (LBVaV virus standard, Prime Diagnostics, The Netherlands), respectively.

Image conversion of the scanned prototypic membrane resulted in an (X, Y, Z) 663 × 458 × 3, 3D matrix of a total of 303,654 pixels, each having unique X-Y coordinates and a particular R, G, B triple integer combination defining its color. After pixel selection, the prototypic image dataset comprised 65,310 observations (rows) and five variables (columns): the pixel color attributes, “Red” (R), “Green” (G), and “Blue” (B), “Sample dilution” and “Infection status.” In turn, the “Sample dilution” variable included six levels (i.e., the undiluted state and the five tested dilutions), whilst the “Infection status” variable included two levels, indicating the presence or absence of the pathogen. The total number of pixels of positive and negative samples were 14,907 and 50,403, respectively. The relative frequencies of pixels of the various categories of the prototypic dataset according to their infection status are presented in ***Table 1***. The pixel information of the samples included in the prototypic DE image was harnessed to train the classifier algorithm.

**Table 1.**
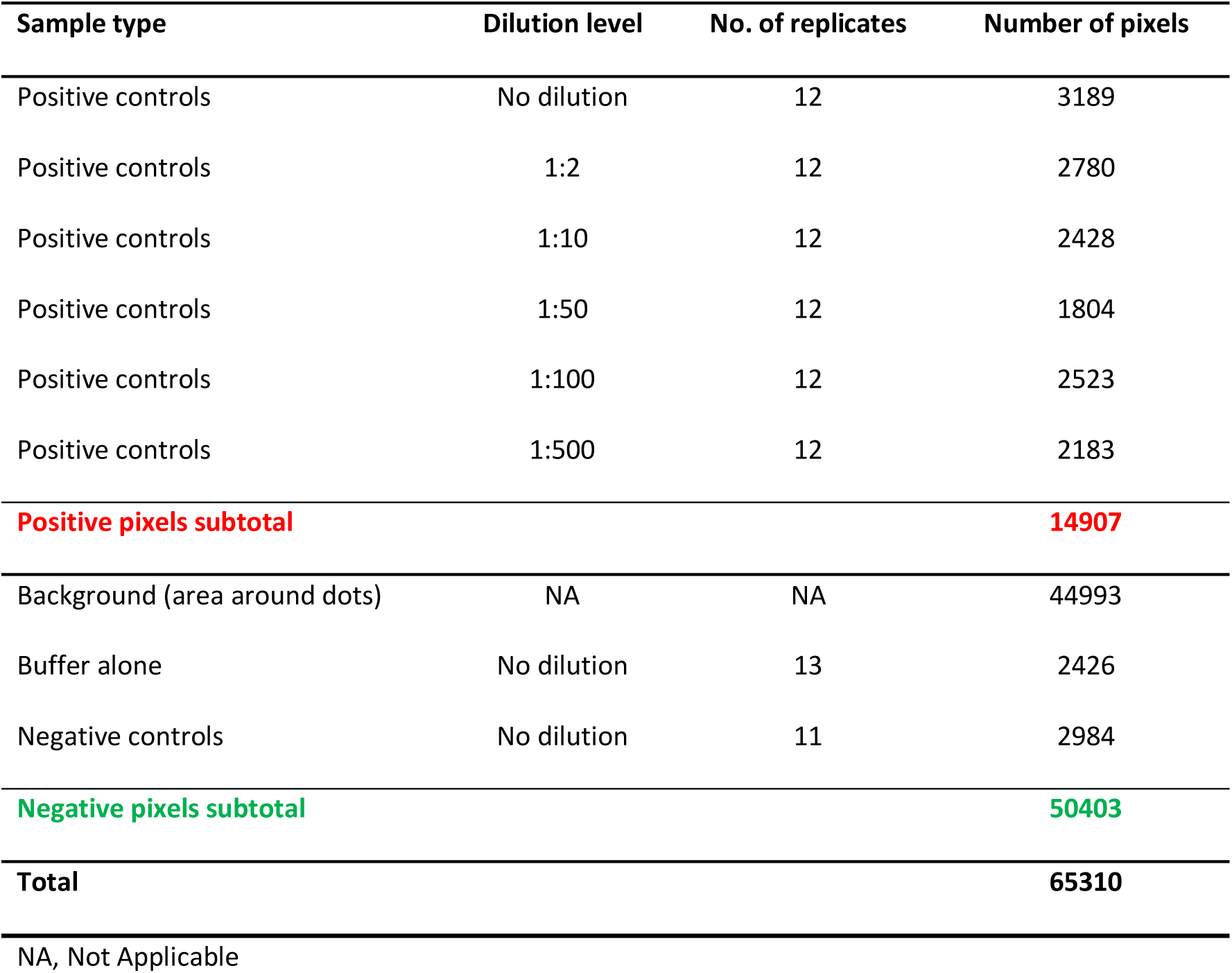
Relative frequencies of pixel categories of the prototypic dataset by infection status.

| Sample type | Dilution level | No. of replicates | Number of pixels |
| --- | --- | --- | --- |
| Positive controls | No dilution | 12 | 3189 |
| Positive controls | 1:2 | 12 | 2780 |
| Positive controls | 1:10 | 12 | 2428 |
| Positive controls | 1:50 | 12 | 1804 |
| Positive controls | 1:100 | 12 | 2523 |
| Positive controls | 1:500 | 12 | 2183 |
| **Positive pixels subtotal** |  |  | **14907** |
| Background (area around dots) | NA | NA | 44993 |
| Buffer alone | No dilution | 13 | 2426 |
| Negative controls | No dilution | 11 | 2984 |
| **Negative pixels subtotal** |  |  | **50403** |
| **Total** |  |  | **65310** |
| NA, Not Applicable |  |  |  |

### Image pixel attributes of positive *vs*. negative samples

Boxplot analysis of all pixels of samples in the prototypic DE dataset revealed unique RGB patterns by infection status (***Figure 3***). In positive controls, the pixel intensity of Green was consistently lower than that of the Red or Blue, whereas in negative controls the intensity of Green was equal to or greater than the Blue. In the buffer alone control, the pixel intensity of Green was also slightly lower than the Blue or Red. Yet, the distributions of the Green and Blue colors of the background area (around the dots) category were similar (and lower than the Red) and their medians tended to be equal. These patterns were preserved when considering one pixel-wide lines traversing each dot of the blot at a specific Y coordinate along the corresponding X coordinates (***Figure 4***). Blue and Green displayed the lowest pixel intensity in negative and positive controls, respectively, in all dilutions. In the highest dilution (i.e. 1:500), although the color intensity was very low, the pattern analogies were maintained. Taken together, these results suggest that pixels of dots of samples in images of DE readouts contain pieces of specifically discriminative information that can be exploited for model training and classification.

**Figure 3.**
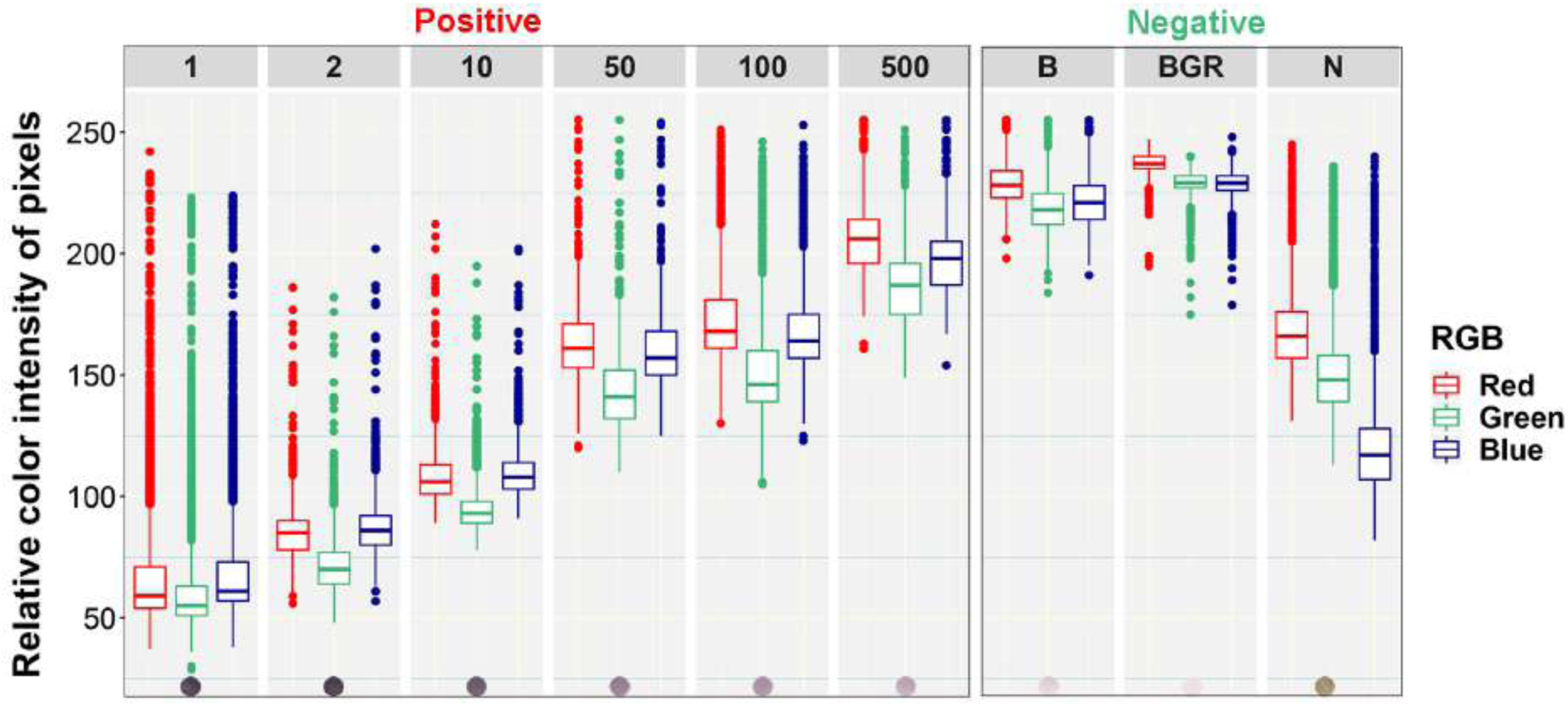
Box plots of relative color intensity of prototypic blot pixels of different categories. Numbers above the plots indicate the undiluted status (1) or dilutions of the positive control (1:2, 1:10, 1:50, 1:100, and 1:500), while letters indicate the Buffer alone (B), background area around the dots (BGR), and Negative (N) controls.

**Figure 4.**
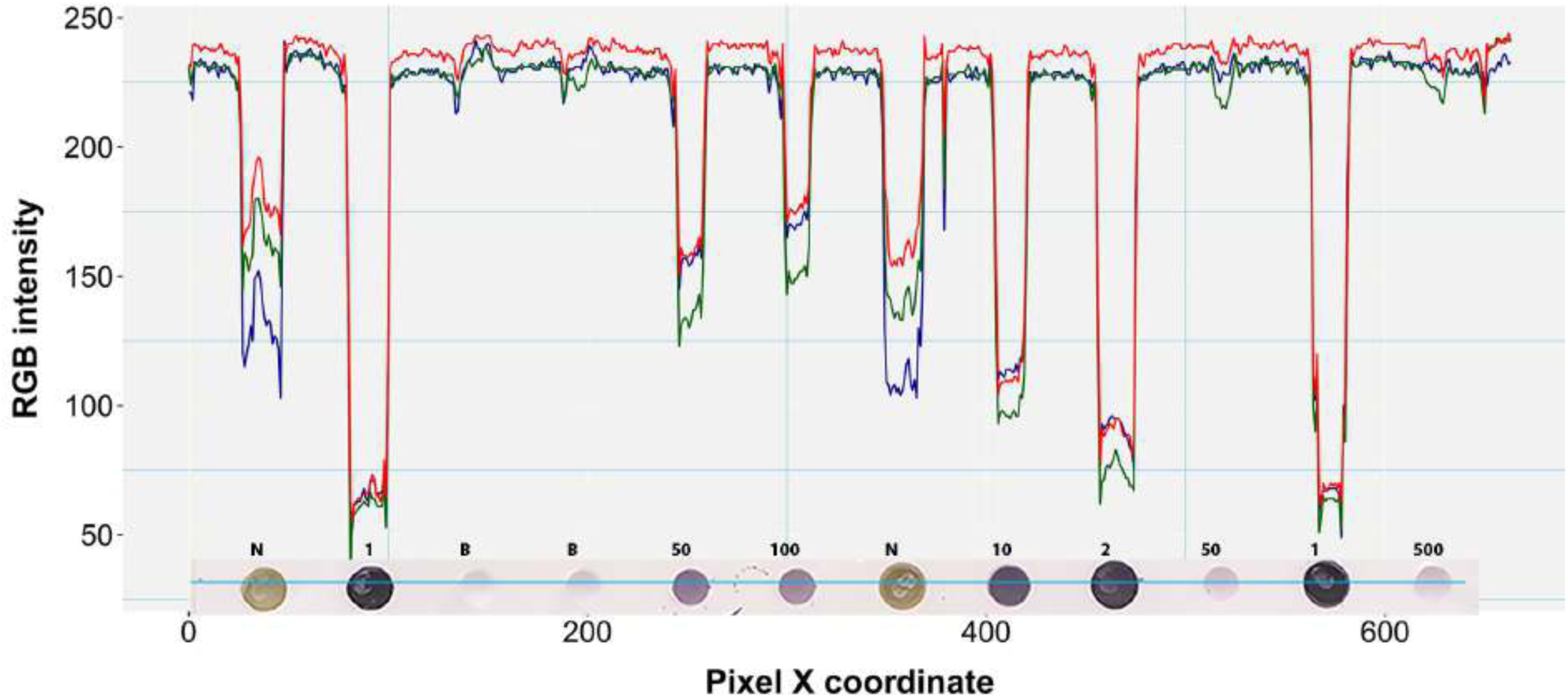
RGB pixel intensity *vs*. dot coloration patterns in a perceived one pixel-wide line. A four pixel-wide line (∼6 pixels per mm) is actually shown here traversing the fourth row of the prototypic blot image at a representative point of the Y-axis (Y=204). Each dot on the blot is a raster or bitmap image of approximately 200 pixels. Numbers above the dots denote the undiluted status (1) or dilution factor of positive samples. N and B denote negative and buffer controls, respectively.

### The RGB model displayed the best performance characteristics for infection status prediction

Among candidate models for infection status prediction, those with B, R and G as single predictors were dismissed for having the highest Akaike information criterion (AIC) values (means ± SEM = 29,592.0 ± 12.69, 21,554.0 ± 11.62 and 20,449.1 ± 11.42, respectively). The R+G combination also had a similarly increased AIC value (19,548.1 ± 11.52), while the AIC of R+B was still elevated, but at about a third of the R+G value (6,263.0 ± 9.42). Although the AIC value of the G+B pairwise combination was further reduced (2,395.0 ± 9.40), it was still larger compared to that of the model constructed using all predictors combined; R+G+B had the lowest AIC (2,382.2 ± 7.05) and was thus selected for training the classifier algorithm. Using the selected model, the R, G, B variable coefficients ± SEM were all found to be highly significant (p<10^−8^) and estimated at β_1_= 0.08 ± 0.015, β_2_= −0.72 ± 0.017, β_3_= 0.54 ± 0.014, respectively, implying a minor contribution of the R variable to positive pixel predictability and a substantial but reciprocal effect of the G and B variables.

### The cutoff (positive/negative threshold) of choice for diagnosis

A comparison of the TP and TN pixel occurrences of the test subset to those predicted by the trained classifier algorithm provided an initial validation of our approach. ***Table 2*** displays the corresponding confusion matrices at five representative cutoff values (0.050, 0.135, 0.500, 0.800 and 0.950). All three calculated diagnostic parameters (accuracy, sensitivity and specificity) were very high (>0.94) across these cutoffs. As expected, when cutoff values increased, TP (sensitivity) and FP (false alarms) decreased, while TN (specificity) and FN (missing values) increased. The relative cost of misclassifying pixels at more or less stringent cutoff values becomes tangible in ***Figure 5*** that shows the overlapping distributions of negative and positive pixels of the test subset of the prototypic image. Yielding the lowest FP and FN rates, the 0.135 cutoff appeared to offer the best trade-off among the various diagnostic performance parameters.

**Table 2.**
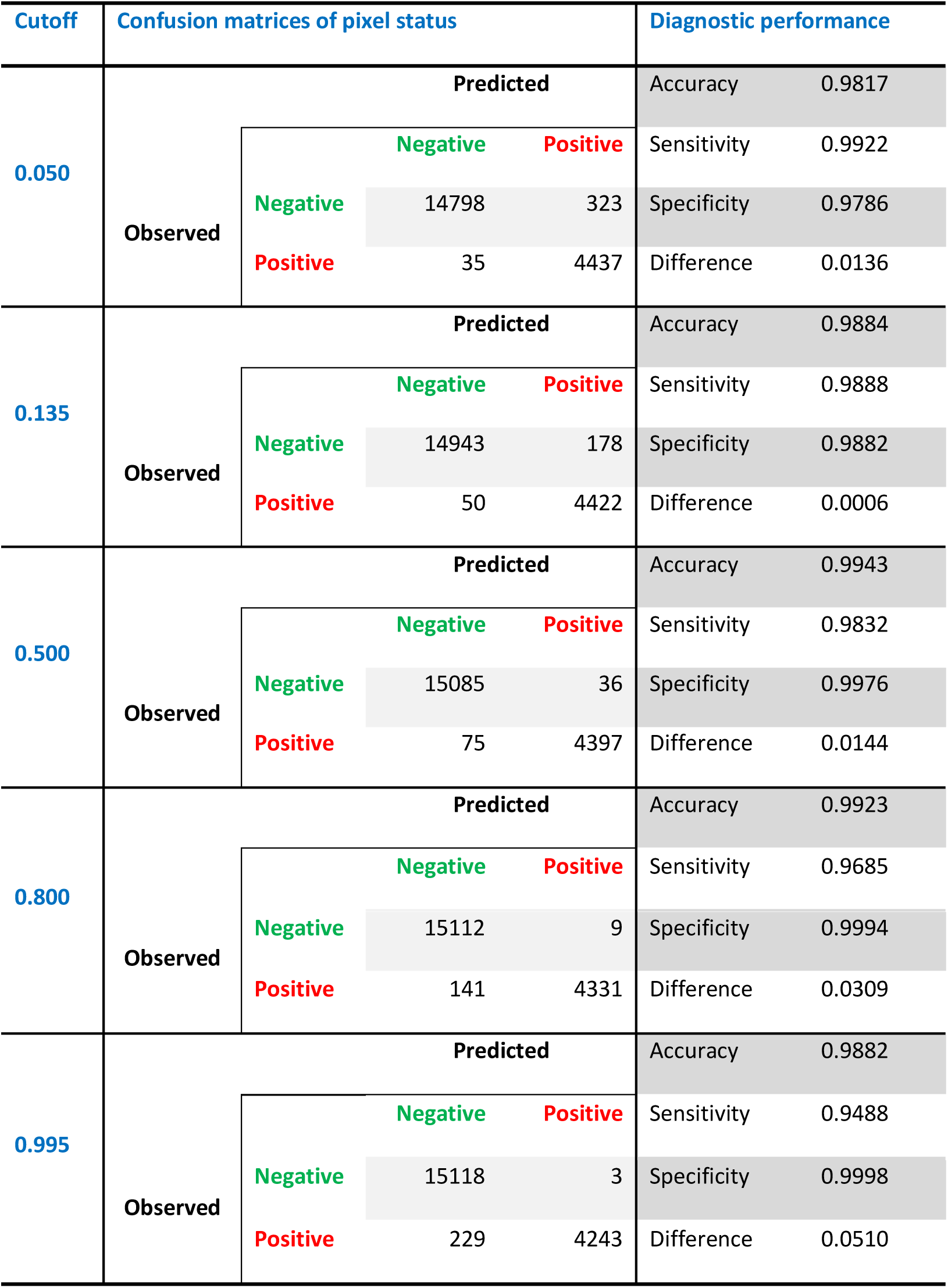
Comparison of diagnostic performance indicators at representative cutoffs generated from the respective confusion matrices of observed against predicted by the trained classifier algorithm pixel status in the test subset (n= 19,593).

**Figure 5.**
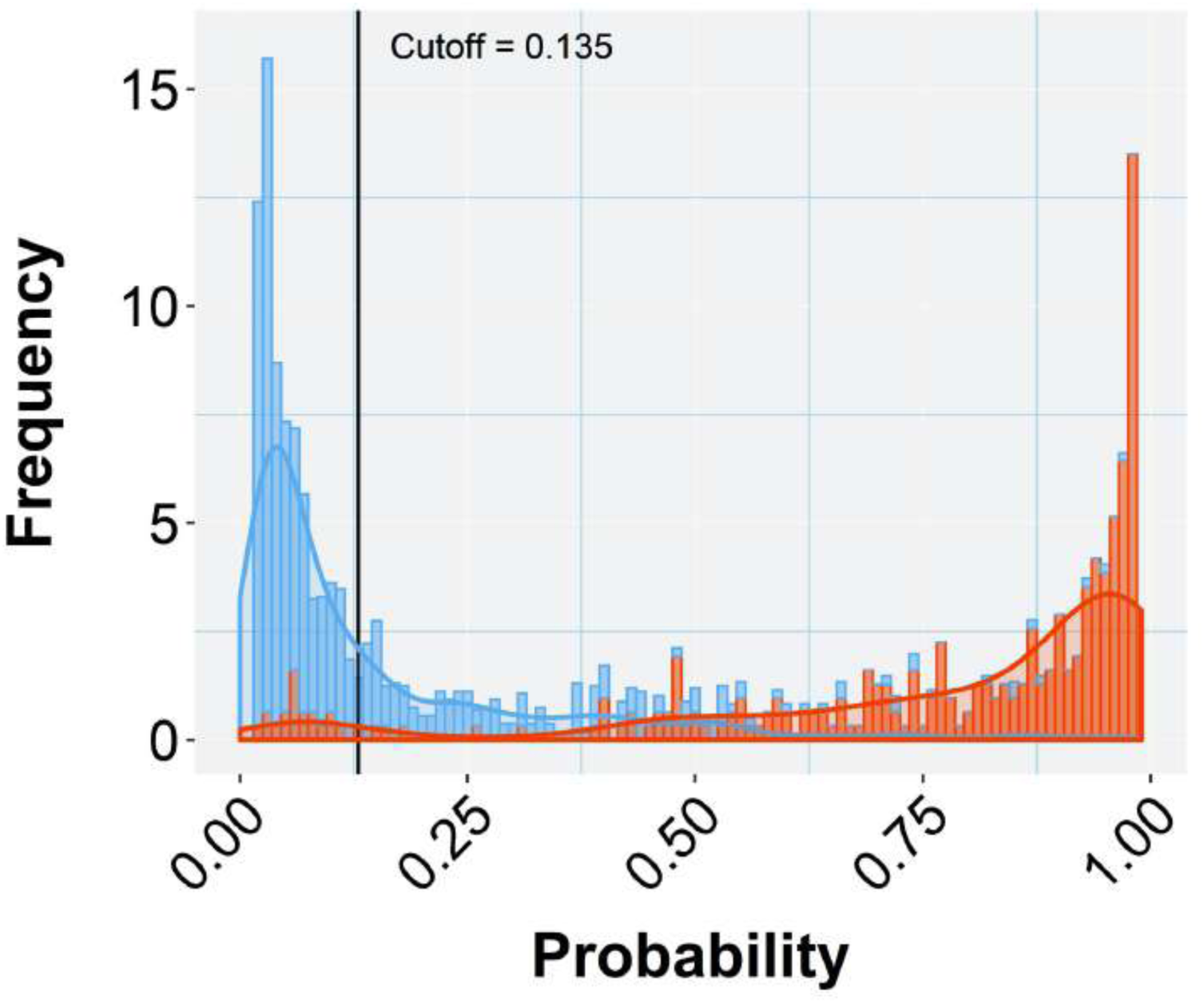
Histogram of stacked bars and the corresponding density curves showing the frequencies and density of classified pixels of the test subset of the prototypic image as negative (blue) or positive (orange). The length of each bar is proportional to the number of classified pixels at each probability class (defined by the bar width). The picture zooms at the range of probabilities between 0.05 and 0.99, concealing the high proportions of negatives (near zero) and positives (near one) to emphasize the overlapping between misclassified pixels. The black vertical line crossing at the selected cutoff (0.135 in this case) divides the plot into two halves that represent negative (below the cutoff) and positive (above the cutoff) predicted pixels. As the vertical line moves to the right (increasing cutoff), more TN are collected (blue bars to the right) at the cost of considering some orange bars (TP) as negatives (TN or specificity increase). When the line moves to the left (decreasing cutoff) more TP (orange bars) are detected at the cost of considering some blue bars (TN) as orange (positives) (TP or sensitivity increase). Selecting the appropriate cutoff for each diagnostic case is unavoidably a compromise that minimizes the effects of the least important error type.

The chosen model with all three-color predictors displayed the best performance indicators at the 0.135 ± 0.0022 cutoff, which was the crossing point between sensitivity (0.9888 ± 0.00009) and specificity (0.9882 ± 0.00009) across the entire cutoff range (***Figure 6***). At this point the two variables differed the least (Difference = 0.0006, ***Table 2***) and consequently, FP and FN rates were at their nadir. Accuracy was also superior for the three-predictor model at this cutoff (0.9884 ± 0.00009) and inaccuracy was about 0.0116 ± 0.00009, which translates to about 230 faulty pixels out of the 19593 pixels of the test subset. The excellent performance of the RGB model was also evidenced by the constructed ROC curve (***Figure 7***), which revealed an Area Under the Curve (AUC) value very closely approximating unity (0.9972 ± 0.00006). The optimal cutoff of 0.135 corresponds to a 98.9% success rate of TP and TN (1-0.0112) predictions.

**Figure 6.**
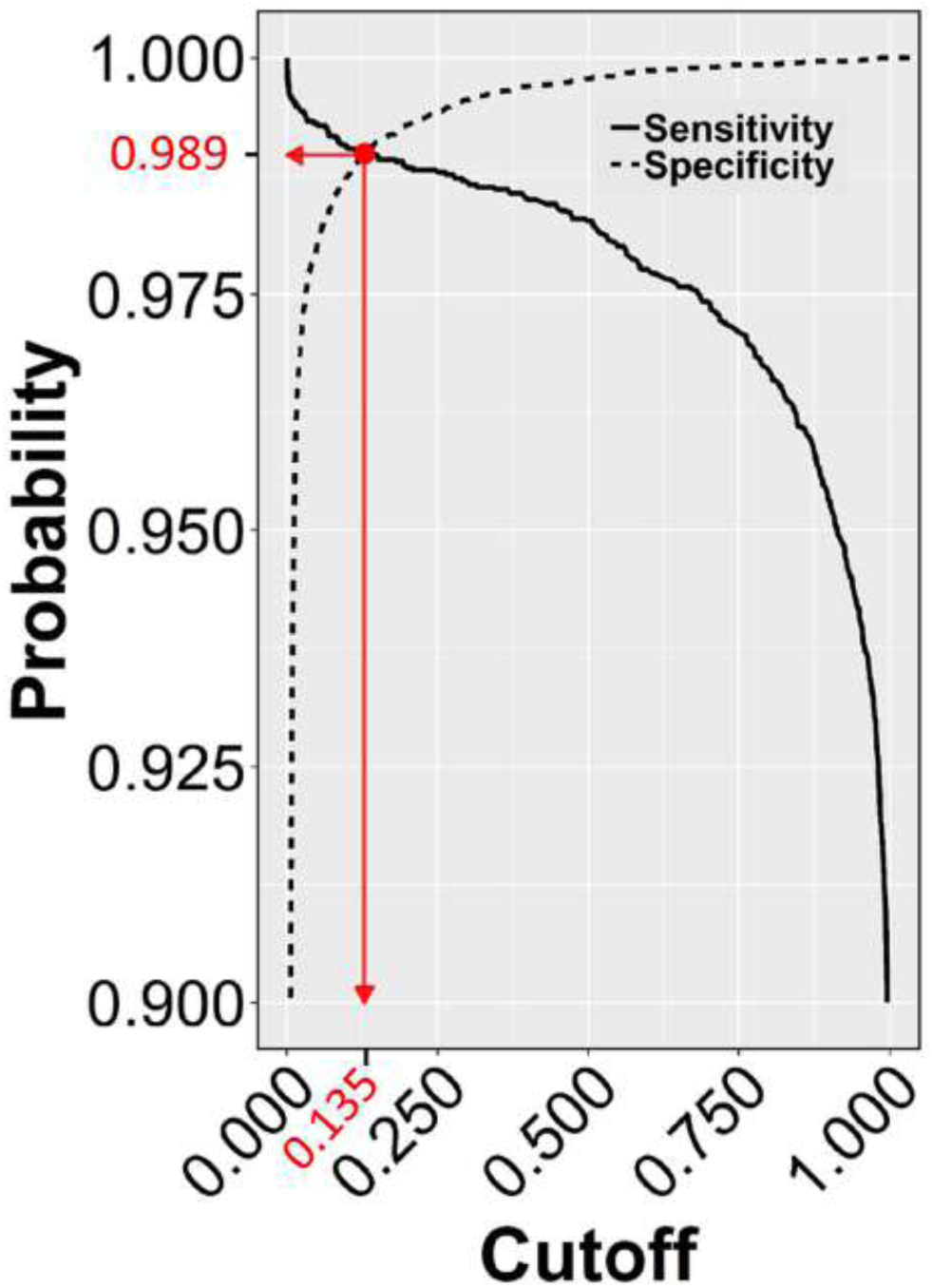
Sensitivity and Specificity of the test dataset across the entire cutoff range (0 – 1). The crossing point (red dot) of the two curves indicates the selected cutoff value (0.135), at the zenith of the TP (0.9888) and TN rates (0.9882), which corresponds to the nadir of FP and FN rates (1-0.9882=0.0118 and 1-0.9888=0.0112, respectively).

**Figure 7.**
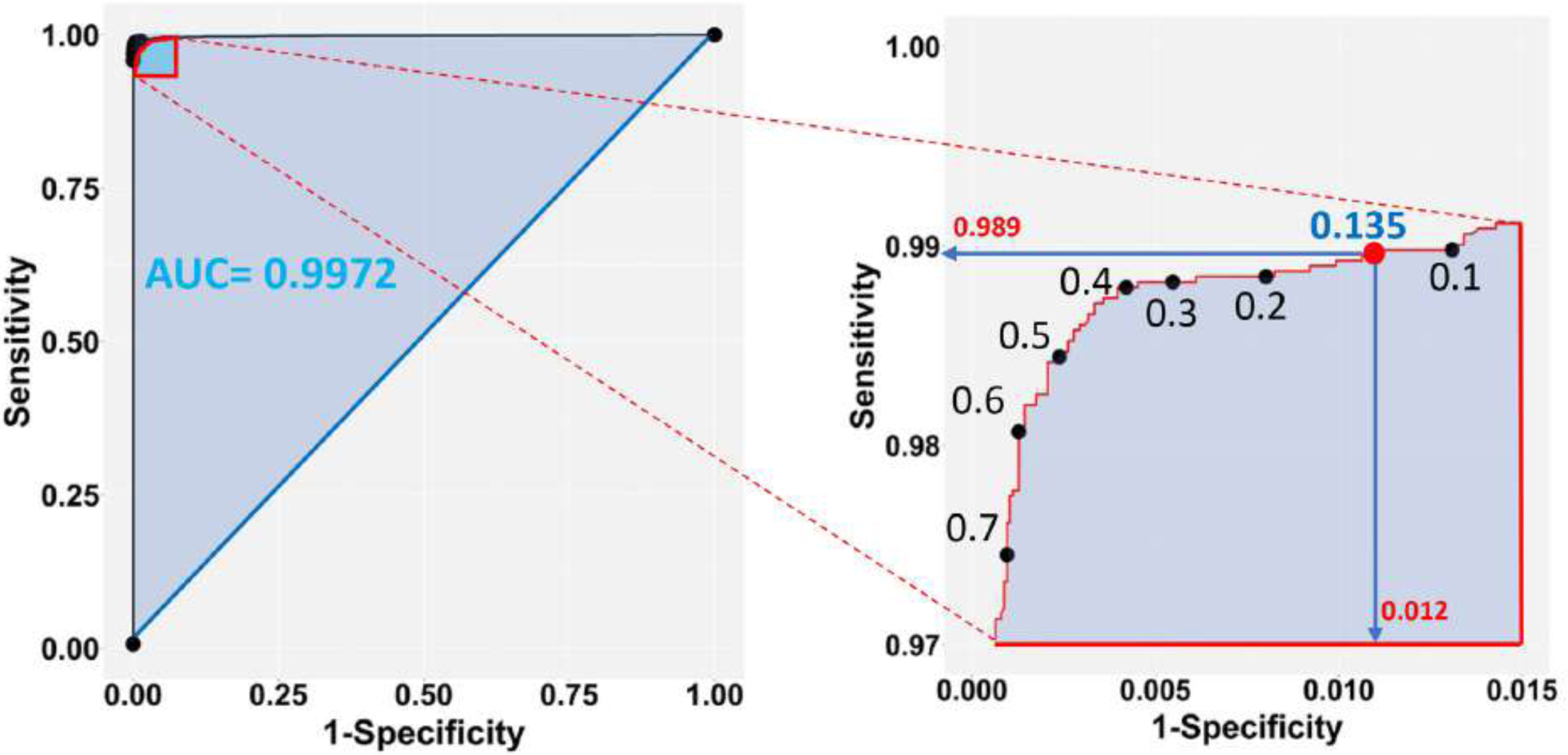
Sensitivity *vs*. 1-Specificity (FP) plot of the test dataset (left panel), zooming into the top left-hand corner of the plot region (right panel). The blue shaded part of the plot in the left shows the Area Under the Curve (AUC). The selected cutoff of 0.135 (red dot) corresponds to a 98.9% success rate of TP and 98.8 % (1-0.0118) TN predictions.

### Unambiguous dot classification and infection status prediction of unknown samples

We applied our method to test plants suspected for *Lettuce big-vein associated virus* (LBVaV) infection. On the original blot (***Figure 8-I***), dots of negative and positive controls displayed the light olive-green and purple color, respectively, as in the prototypic DE output (***Figure 2***). Therefore, the olive-green dots of unknown samples (e.g. C5:D5, C7:D7, E10:F10, and E12:F12, ***Figure 8-I***) resembling negative controls (G9:H9 to G12:H12) could be classified rather easily as virus-free. However, most dots of unknown samples displayed different shades of purple, ranging from cyclamen to deep violet. Some dots were lighter (e.g. C4:D4 and E4:F4), comparable (e.g. A12:B12 and C12:D12), or darker (e.g. A1:B1, A2:B2, E8:F8 and E9:F9) than the positive controls, rendering infection status determination by eye-based comparison uncertain. Application of our trained and validated algorithm cleared up the confusion. After matrix conversion and pixel probability prediction, the DE image of unknown samples was reconstituted at various cutoffs (***Figure 8, II-VI***), revealing truly positive samples with a very high probability of success. Positive controls produced a robust signal throughout the range of examined cutoffs, while negative controls did not produce a signal at any considered cutoff value, as was the case for the Buffer alone negative controls.

**Figure 8.**
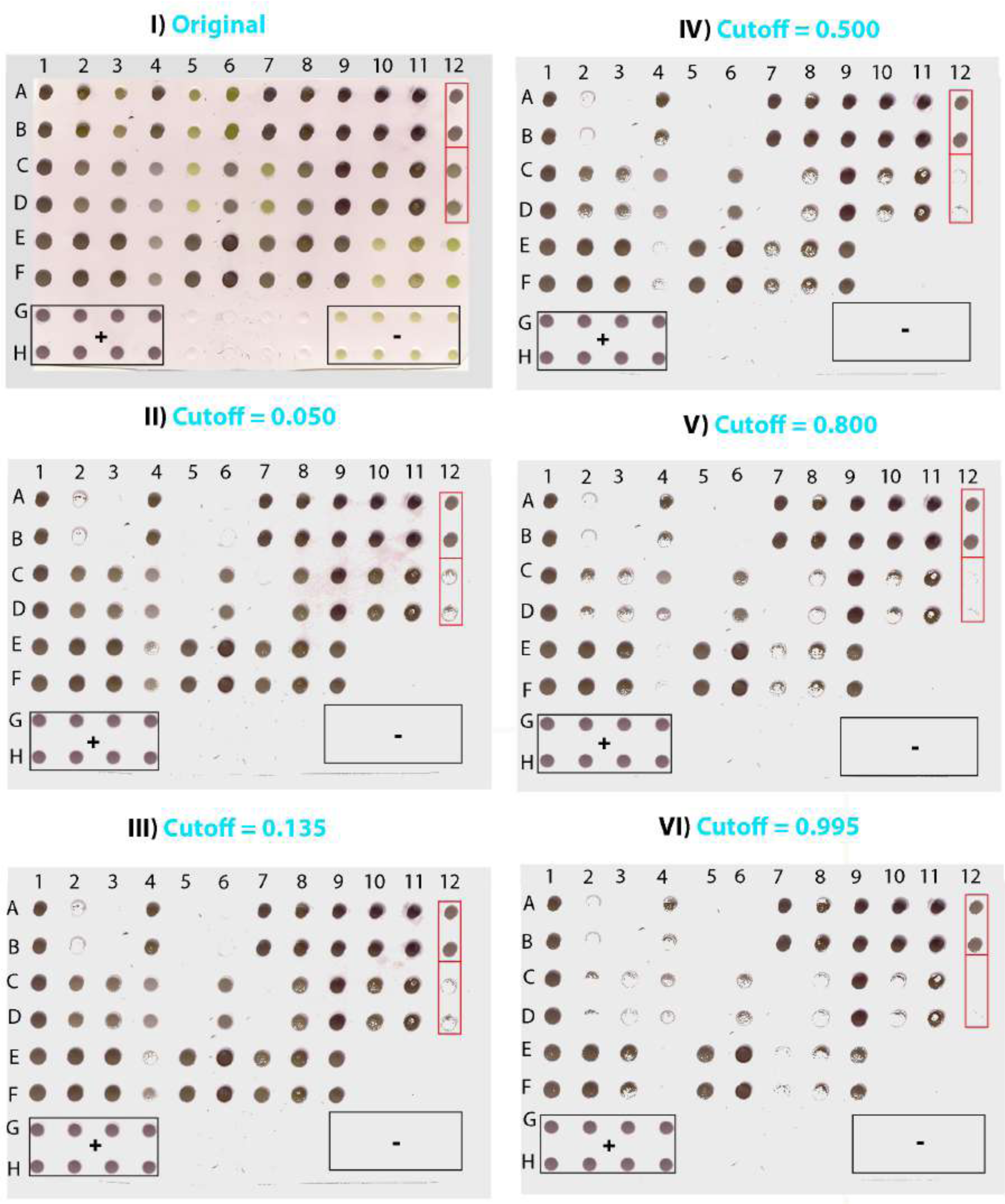
Scanned image of a DE output of lettuce samples suspected for LBVaV infection (I) and the corresponding images reconstituted from pixels applying cutoffs of 0.050 (II) 0.135 (III), 0.500 (IV), 0.800 (V) and 0.995 (VI). Duplicates of 36 samples of unknown infection status were loaded in [(A-F):(1-12)]. Lanes G and H contain duplicates of positive (+) controls (1-4), Buffer only (5-8), and negative (-) controls (9-12). Framed in red are two of the sets of dots of ambiguous diagnosis that appear similar in the original DE output (I). In the reconstituted images, however, where pixel information of dots has been harnessed, it becomes clear that dots A12:B12 are positive and C12:D12 negative.

***Table 3*** shows the relative preponderance by increasing cutoff values of positive versus negative pixels of unknown samples classified by the trained algorithm as well as the corresponding dots calculated from pixels by accepting a mean value of 237 pixels per dot, or actually detected in the reconstituted images of the DE output of unknown samples. As the cutoff increased, so did the negative pixels and the corresponding dots, either calculated from the pixels or actually detected (the two categories differed less than 3%, i.e., by a maximum of two dots out of the 72 of unknown samples). The opposite trend was noted among positively detected pixels and the respective dots, the numbers of both of which decreased as the cutoff increased.

**Table 3.**
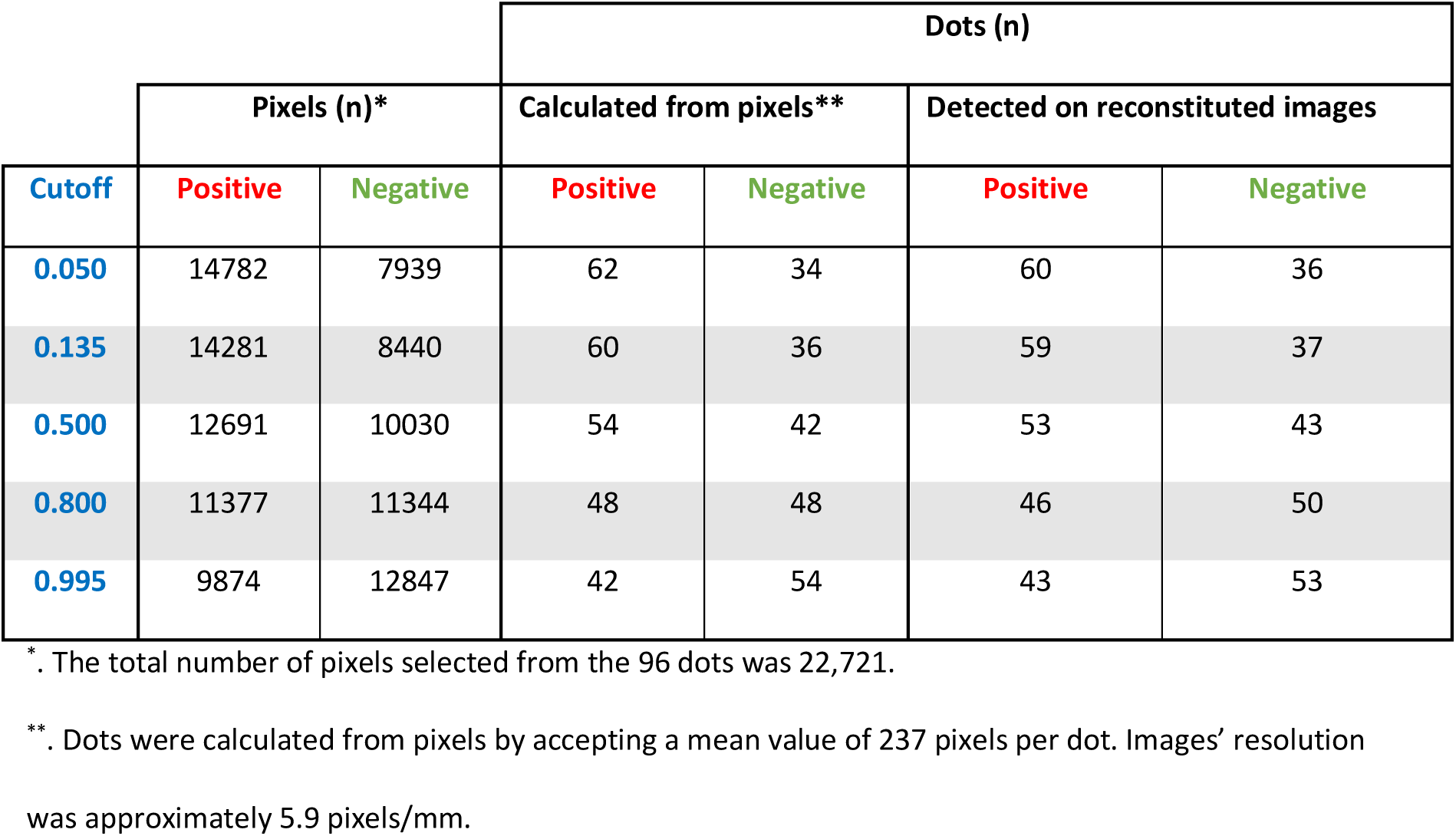
Relative numbers of predicted positive and negative pixels of dots of the DE output of unknown samples and the corresponding numbers of dots calculated from pixels or counted in the reconstituted images by increasing cutoff values (related to ***Figure 8***).

At the very low cutoff of 0.050, which corresponds to low specificity and low FN rates (***Figure 8-II***), all dots suspected to be negative due to their color hue (light olive-green in ***Figure 8-I***) were classified as such. Nevertheless, some dots that were practically indistinguishable by eye from positively classified pairs on the original blot also appeared to be negative (e.g. A2:B2 and C12:D12 *vs*. A1:B1 and A12:B12, ***Figure 8, I & II***, respectively). One of the duplicate dots that gave a weak positive signal at the 0.050 cutoff (E4) appeared negative at the 0.135 cutoff, which corresponds to the lowest FP and FN rates (***Figure 8-III***). At the more elevated cutoff of 0.500 (***Figure 8-IV***), some dots that appeared positive at the 0.135 cutoff were found to be negative (e.g. E4:F4), whereas some others appeared sparsely populated by positive pixels (e.g. C8:D8, C10:D10, and E7:F7).

These findings are consistent with a trend towards increasing FN rates with higher cutoff values as also suggested by the pixel *vs*. dot data presented in ***Table 3***. Thus, at the 0.800 cutoff more positive pixels were lost from the sparsely populated dots detected at the previous cutoff; some of these dots appeared to be negative (e.g. C8:D8), whereas some others were still positive (e.g. C3:D3, E8:F8) (***Figure 8-V***). At the highest cutoff examined (0.995), where sensitivity and FP rates are expected to be even lower, dots yielding positive signals had the highest probability of being TP (***Figure 8-VI***). Here, only the darkest dots maintained their compactness, whilst most sparsely populated dots detected at the previous cutoff appeared negative (e.g. C3:D3 and E8:F8).

To test the versatility of the method, we examined an additional DE output loaded with representative dilutions of *Mirafiori lettuce big vein virus* (MilBVV), a virus unrelated to the training procedure. All TP and TN samples were correctly predicted as such in the reconstituted image of the scanned DE output, corroborating the versatility of the method (*Figure 9*).

**Figure 9.**
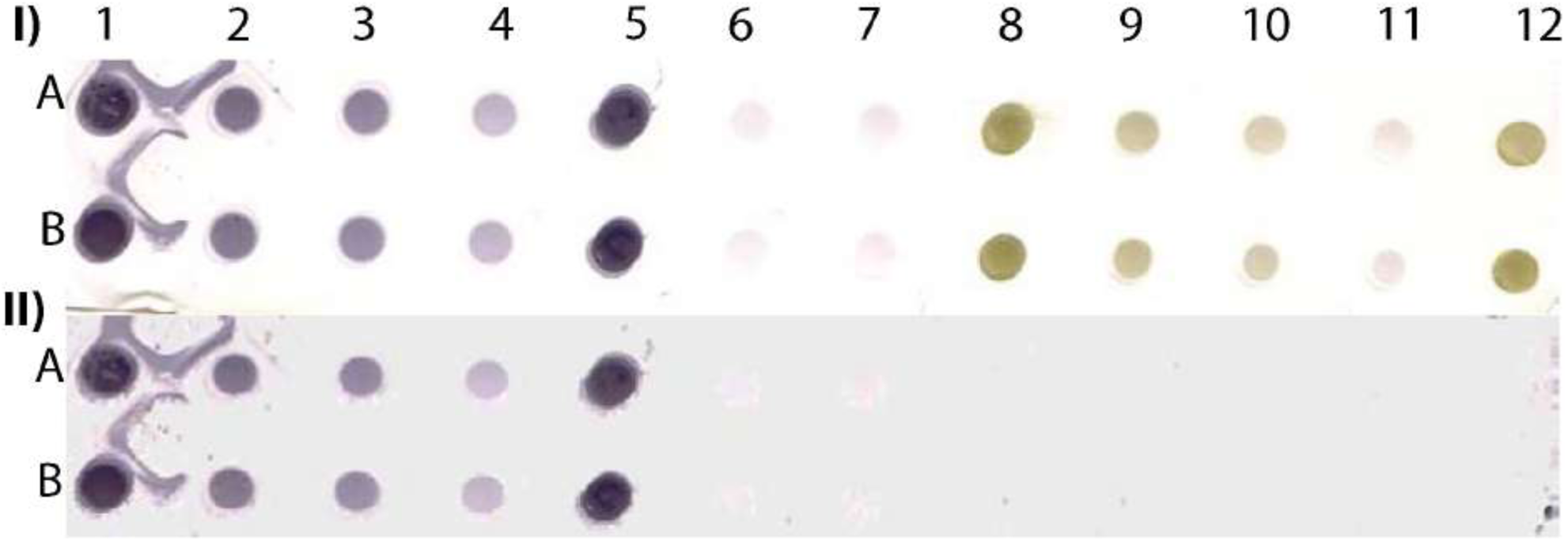
Scanned image of a DE output of positive and negative controls of MilBVV (I) and reconstituted image (II) made by pixels with predicted probabilities above the 0.135 cutoff based on the LBVaV-trained model. Lanes A and B show duplicates of undiluted (1 and 5), and 1:5 (2) 1:10 (3) and 1:50 (4) dilutions of the positive control of MilBVV. Dots 6 and 7 are buffer only controls. Dots 8 to 12 contain the negative control of MilBVV undiluted (8,12), and at 1:5 (9), 1:10 (10), and 1:50 (11) dilutions. The translation of unique antigens, of MilBVV here, to the universal basic color language allows for its detection by the algorithm (model) we built, even though it was trained on the basis of a different virus, LBVaV.

## Discussion

Diagnostic procedures for pathogens differ depending, inter alia, on the physiology of the infected organism. In animal species including humans, diagnostic methods, such as nucleic acid testing, allow for the direct detection of the microorganism causing the infection, typically after an amplification step using, preferentially but not exclusively, the polymerase chain reaction (PCR), whilst indirect methods, such as serological assays, allow for the detection of antibodies against microorganism antigens. Pathogen detection in plants that lack an immune system do rely on the proper detection of the pathogen, but with a couple of notable exceptions: the amplification of specific targets through PCR is often proven difficult due to the presence of inhibitors in plant cells (***Dorn et al., 1999; Lacroix et al., 2016; Lardeux et al., 2016; Rački et al., 2014; Schrader et al., 2012; Suther and Moore, 2019***) whereas, the role of the detected pathogen component is rather obscure at least from the etiological point of view. In the case of viral plant pathogens, for instance, the capsid protein subunits or the virus particles containing the viral genome may be detected simultaneously (***Manoussopoulos and Tsagris, 2015***), indicating the establishment of the virus in its host. ELISA and its inexpensive and even more sensitive, solid-state alternative DE are the methods of choice for quick and reliable pathogen diagnosis in both analytical and epidemiological surveys in plants (***Henry Sum et al., 2017***). DE has also been used for disease diagnosis in humans (***Rodkvamtook et al., 2015; Subramanian et al., 2016***) and animals (***Fisa et al., 1997***), but mostly in resource-poor settings. The major obstacle limiting the generalized applicability of the method is uncertainties in DE output evaluation. In this work, we present an innovative, flexible and reliable unbiased approach through which unique pathogen antigens are translated into the universal, basic color language of pixels (RGB) that may be used for unambiguous DE output evaluation.

The proposed method relies on supervised machine learning based on a logit (logistic) function that is used for prediction purposes. Machine learning is, in the broader sense, an artificial intelligence procedure that enables a computer (machine) to identify patterns in big datasets on the basis of previous training on known data (***Alpaydin, 2014***). Classification is a type of supervised machine learning in which the learning algorithm (model) is trained on a set of rules that enable the correct discrimination of items into categories (***Raschka and Mirjalili, 2017***). Depending on their dimensions and resolution, raster images consist of thousands to millions of pixels containing information that may be exploited for classification purposes. Raster images can be obtained easily via a scanner or a similar device and, in turn, they can be readily converted into 3-D matrices and datasets, holding the pixel RGB attributes and X-Y position-specifying coordinates in the image. Our analysis showed that the RGB values of dots of DE outputs remained constant at different antigen dilutions, providing discriminating information suitable for training in machine learning applications. Differences in RGB patterns persisted even at the highest sample dilutions. Although not easily discriminable by eye evaluation, samples at high dilutions had distinguishable RGB patterns from those of buffer control or the background. Thus, DE blots scanned to raster images may be used in machine training not only for discriminating negative from positive samples but also for distinguishing between low concentration antigens and background, a task currently challenging in serodiagnostic techniques.

To the best of our knowledge, machine learning has not been exploited for DE output evaluation. The method proposed herein is simple and requires no expensive accessories to run; a computer and a common portable/handheld scanner are all that is needed. The innovation at the heart of the procedure lies on the exploitation of image pixels, which convey invaluable information ideally suited for machine training, thereby allowing for predictions to be made for unknown pixel status with measurable probabilities of success. Concerning previous work in the field, only a few attempts have been made mainly based on deep learning with most of them serving to complement other procedures. Thus, a deep learning approach has been employed to predict positives in ELISA microplates (***Nath et al., 2018***) based on the training of an artificial neural network with microplate images of known sample status and the application of the trained algorithm on microplate images of unknown samples. In another report, machine learning was used as an additional step in a procedure used to evaluate transformed ELISA microplate images of a Cellphone-Based Hand-Held Microplate Reader (***Berg et al., 2015***). In a more recent work, an artificial intelligence system was presented based on supervised deep learning that outperformed human expertise in the evaluation of mammograms for breast cancer prediction, demonstrating the great promise that such systems hold for diagnostic applications (***McKinney et al., 2020***).

ELISA is a widely used method for serodiagnosis of many important human, animal and plant pathogens. One drawback of the method is the need for sophisticated and expensive laboratory equipment that increases the per test cost. Another, perhaps more important disadvantage, is the lack of assessment of FP and FN rates that limits its application solely to high accepted cutoff (positive/negative threshold) values that correspond to increased FN, unavoidably leading to the loss of TP. Indeed, lacking a reliable method for cutoff estimation, the general rule of thumb for in-house ELISA microplate evaluation is the acceptance as positives only of those samples having OD values either two or three times the mean of negative controls (***Crowther, 2009; Lardeux et al., 2016***), resulting in elevated FN rates. The situation is not less problematic for commercially available ELISA kits, where manufacturers often claim comparable sensitivity and specificity to other commercially available - and even licensed - ELISA kits, without elaborating further on the actual diagnostic performance of the assays.

However, serodiagnostic methods are ever important in both clinical, veterinary and phytopathological practices, complementing the so-called molecular methods, exemplified by, but not restricted to, hybridization and PCR, or the recently developed advanced sequencing methods known as next (NGS) and third generation (TGS) sequencing. Although extremely useful as an ultra-sensitive and highly specific diagnostic tool, PCR, in its various formats, needs laborious optimization and stringent conditions to avoid the risk of contamination, it is costly and time-consuming with rather low throughput since nucleic acid isolation and possibly reverse transcription steps have to precede it. The newly developed NGS and TGS, on the other hand, are highly informative for unknown or candidate pathogens, but are still very expensive for diagnostic purposes and they tend to yield results that are difficult to interpret, particularly in the field of etiological host-pathogen interactions. Compared to these approaches, serological methods are cheaper, require less sophisticated equipment and are better suited for large surveys for the detection of known pathogens in established host-pathogen relationships. Serological monitoring is essential, in particular, to the design and evaluation of effective vaccination programs. DE inherits all advantages of the maternal ELISA application and overcomes most of its disadvantages pertaining to cost and the need for specialized equipment, rendering it an attractive alternative for serodiagnosis in epidemiologic research once the problem of output evaluation is resolved.

Besides fully addressing in an objective and reliable manner the issue of output evaluation, a major advantage of our method is that it allows for cutoff selection, thus enabling decisions to be made on FP and FN acceptance rates suited to the diagnostic question at hand; furthermore, estimated probability values are calculated for TP, TN, accuracy and all types of errors. The trained model showed outstanding performance with high accuracy and very low FP and FN rates. The accepted values of FP and FN could be easily selected through the generated ROC curve. These features potentially enhance assay reproducibility by eliminating signal variability among experiments arising from fluctuations in pathogen antigenicity levels or pathogen-antibody and/or antibody-conjugate affinities. Our findings demonstrate the unambiguous recognition of TP and TN dots in DE outputs, with certain probabilities of success, a feature missing from serodiagnostic applications. Another advantage of the method is the short time it requires since no image preprocessing is necessary and only a few tens of seconds, depending on the size of the image, are needed to scan the processed blot and reconstitute the image from pixels above the selected threshold.

Importantly, the method is both antigen- and control-independent. As shown in the proof-of-concept experiment, the trained by the LBVaV antigen algorithm successfully detected all tested dilutions of the MilBVV antigen. The two lettuce big-vein disease-associated viruses belong to different classes and, as such, they have completely different molecular and particle structural properties. This universality stems from the fact that there is no relationship between the kind of antigen and the color produced; therefore, the method is highly versatile and appropriate for the serodiagnosis of antigens of any microorganism. The algorithm classifies image pixels based on their color intensity, rendering retraining and the inclusion of controls with each run unnecessary, so long as the same chromogenic substrate is used. The DE control-independence offers a great advantage over ELISA, where output evaluation is chiefly dependent on positive and negative controls, limiting the number of samples that can be tested simultaneously and increasing cost.

The method can be further improved by training the algorithm with pixels from many more prototypic images, covering all possible dot colorations from the full range of dilutions, with emphasis on the lower end of the spectrum where positive signals turn blurry and indistinguishable from the background. In theory, the training dataset could be developed by code that calculates the entire range of RGB values specific for the substrate spectrum of color intensities, thereby avoiding prototypic blot preparations which cannot easily cover the whole spectrum of colors. Both the incorporation of more exploratory variables (e.g. dilution levels) in the training model and the employment of different models, such as classification trees or random forest algorithms, could be also explored. In addition, the whole procedure could be further developed to a web-based application where the input will be the scanned DE image of interest and the output will be the reconstituted image showing the positive and negative dots along with useful statistical metadata.

Application of the described method could contribute to the rapid and reliable diagnosis of common or emerging pathogens in both developed and developing countries since serological methods are probably the most economical and rapidly applicable methods available.

## Materials and methods

DE output evaluation may be reduced to a binary classification problem since test samples can be either infected or not infected with a given pathogen. Machine learning can be used to train a classifier algorithm on known sample attributes to predict the infection status of unknown samples. The following steps are key to this process: (1) prototypic DE dataset construction; (2) model selection with appropriate predictive variables of high discriminative power and supervised training of the classifier algorithm; (3) validation of the trained algorithm; (4) ROC curve analysis and cutoff selection; (5) infection status prediction of unknown samples. An overview of the method is presented in ***Figure 1***, while each step is described in the sections that follow.

### Step 1: Prototypic DE dataset construction

The prototypic dataset (***Figure 2***) was constructed from a scanned output image of known positive and negative controls following DE undertaken according to standard protocols using the Bioblot apparatus (Bio-Rad Laboratories, Redmond, WA, USA) and commercially available LBVaV antigens and specific antibodies (Prime Diagnostics, The Netherlands). Volumes of 100 μl of the following types of samples were loaded on the prototypic nitrocellulose membrane: (i) virus standard prepared according to the manufacturer’s instructions (1 OD in ELISA at 405 nm in 30 minutes, Prime Diagnostics, The Netherlands), undiluted or diluted 1:2, 1:10, 1:50, 1:100 and 1:500 in 0.01 M PBS at pH=7.4 (positive controls); (ii) healthy lettuce extracts diluted 1:10 in the same phosphate buffer (negative controls); and (iii) buffer alone (additional negative control for assessing background noise). Positive, negative and buffer controls were randomly distributed on the 96-well apparatus in twelve, eleven and 13 replicates, respectively. A blocking stage [5% skimmed milk in PBS plus 0.05% Tween (PBS-T)] was included after sample loading. The membrane was then processed with the corresponding specific secondary, AP-conjugated antibodies. Washing for five minutes with PBS-T was applied three times between each stage and after color development following incubation with BCIP/NBT substrate for 15-20 minutes. The blot was then dried on Wittman paper and scanned as a TIFF color image at a resolution of 150 dpi.

Pixels in the scanned image of the prototypic DE dataset belonged to the following four categories: positive, negative, and buffer or background controls, reflecting the three types of loaded material and the area around them, correspondingly. The positive category was further divided into six subcategories reflecting the undiluted state or the five dilution levels of the virus standard. The prototypic dataset was created by manually selecting regions of each pixel category in the image (i.e., respective dots, but also including representative regions of the background) and storing and processing the X-Y coordinates and RGB information of the pixels using the ImageJ package (***Schneider et al., 2012***).

### Step 2: Model selection and supervised training of the classifier algorithm

The constructed prototypic DE dataset was used for model variable exploration and machine learning. Statistical analysis and training were performed by programming the proper library or function of the “R” statistical program (***R Core Team, 2018***). A multivariable logistic regression model was employed in which the log odds of infection status (i.e., “Yes” or “No”) was the dependent variable and the R, G, B attributes of the pixels were the predictor variables. The generalized linear model had the following form (***Agresti, 2013***):

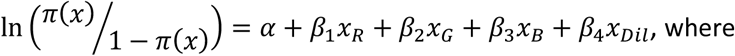

the letters α, β_1_, β_2_, β_3_, β_4_ represent the intercept and corresponding parameters of the x_R_ (Red), x_G_ (Green), x_B_ (Blue), x_Dil_ (Dilution) variables. Successful event (positive pixel) probabilities can be obtained from the model according to the equation:

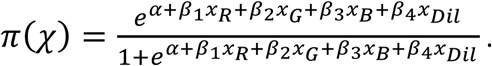

To select the most appropriate predictors, the AIC value of different logit models was considered. Since AIC provides an estimate of the relative amount of information lost by a given model, the less information a model loses, the higher the quality of that model (***Akaike, 1974; Hosmer et al., 2013***). Each model was constructed using a single predictor (i.e., R, G, or B), each of all possible pairwise predictor combinations (i.e., R+G, R+B, or G+B), or all three predictors combined (i.e., R+G+B). In all cases, the ‘Dilution’ category and variable interactions were excluded at this stage for simplicity. Logistic regression was performed using the “glm” function in “R” (***R Core Team, 2018***).

### Step 3: Validation of the trained algorithm

Training was performed using the best fit logistic model and the prototypic subset. To avoid data overfitting (overtraining), an additional cross-validation step was undertaken using two random subsets obtained from the prototypic DE dataset by employing the “caTools” library (***Tuszynski, 2018***) in R (***R Core Team, 2018***): a training subset holding about 70% of pixels and a test subset holding the rest 30% of pixels of the original (prototypic) image (***Figure 2***). For validation, predictions on the “training” and “testing” subsets were made by the “predict” function in “R” based on the logistic training model and the training subset. Confusion matrices were constructed at 0.01 intervals for each randomly created dataset and critical diagnostic parameters, such as the Cutoff, Sensitivity, Specificity, Accuracy and Error, were calculated as follows: Sensitivity was estimated as TP/(TP+FN), Specificity as TN/(TN+FP), Accuracy as (TP+TN)/(TP+TN+FP+FN) and Error as (FP+FN)/(TP+TN+FP+FN). The optimal cutoff was estimated as the probability at which Sensitivity and Specificity differed the least. The above process was repeated 100 times using code written in “R” (***R Core Team, 2018***). For validation purposes, the same diagnostic parameters of each random dataset were obtained using the “performance” function of the “ROCR” library (***Sing et al., 2005***) in “R” (***R Core Team, 2018***).

To obtain 95% confidence intervals (CI) for the mean of each variable, the corresponding dataset containing a random sample (n=100) of each of the parameters was bootstrapped 10,000 times using the “boot” library (***Davison and Hinkley, 1997***) in “R” (***R Core Team, 2018***). To ensure reproducibility and integrity of the used datasets, which could be influenced by stochasticity introduced during the dataset splitting, each of the 100 splitting events was controlled by the R base “set.seed” function, taking incrementally integer arguments from 1 to 100, with each integer corresponding to one splitting event.

### Step 4: ROC curve analysis and cutoff selection

The ROC curve of each dataset was constructed using the “ROCR” library (***Sing et al., 2005***) in “R”. Representative cutoffs and the corresponding Sensitivity and Specificity values were obtained from constructed ROC curves. Optimum cutoff selection and corresponding variables were obtained by finding the minimum absolute difference between “Sensitivity” and “Specificity” that coincided with crossing point of the two variables. AUC was calculated at the selected cutoff of 0.135.

### Step 5: Infection status prediction of unknown samples

To investigate the diagnostic performance of the method and its applicability to real-world situations, we used lettuce plants grown in a field infested with the fungus *Olpidium virulentus*, a known vector of LBVaV (***Roggero et al., 2000***). Sap from lettuce leaves was extracted (1g/10 ml) in phosphate buffer (0.01M, pH=7.4) and centrifuged at low speed (10,000g) for ten minutes. The supernatant was collected and loaded onto nitrocellulose membranes, which were processed with LBVaV-specific, AP-conjugated antibodies and NBT/BCIP substrate as described previously. To further examine the detection performance of the method for an antigen other than that on which the training model was based, we followed the described procedure for a DE blot loaded with representative dilutions of commercially available MilBVV positive and negative controls (Prime Diagnostics, The Netherlands). After color development and washing, dried sheets in each case were scanned into color images and the corresponding matrices with pixel coordinate information and RGB values were obtained by programming the “sys”, “NumPy” and “pandas” libraries in Python ver. 3.5 (***Oliphant, 2007; van der Walt et al., 2011; Wes McKinney, 2010***). Pixel probabilities of the unknown sample datasets were predicted using the “predict” function in “R” (***R Core Team, 2018***) and the trained subset logistic model. DE images of unknown samples were reconstituted at a broad range of cutoffs to investigate their effects on infection status predictions. Pictures were reconstituted by programming the “ggplot2” library (***Wickham, 2016***) in “R” (***R Core Team, 2018***) using pixels falling above the representative cutoffs of 0.05, 0.16, 0.5, 0.8 and 0.995 obtained from the ROC curve or from the described explorative confusion matrices, along with the corresponding pixel X-Y coordinates. The DE image of MiLBVV dilutions was reconstituted at the 0.13 cutoff.

### Predicted pixels’ classification and dot infection status

To further examine the association between predicted pixels’ classification and dot infection status, we manually collected all X-Y coordinates and RGB information of the dots of the unknown samples using ImageJ (***Schneider et al., 2012***). Subsequently, we made predictions using the trained algorithm and calculated the total positively and negatively classified pixels across the range of representative cutoffs, i.e. at 0.050, 0.135, 0.500, 0.800 and 0.995. Then, using the mean number of pixels per dot, we estimated the positive and negative dots corresponding to the reconstituted pictures of each of the examined cutoffs.

## Author contributions

Cleo Anastassopoulou, Visualization, Methodology, Supervision, Project administration, Writing— original draft, Writing—review and editing; Athanasios Tsakris, Resources, Writing—review and editing; George P. Patrinos, Resources, Writing—review and editing; Yiannis Manoussopoulos, Conceptualization, Data curation, Software (Code development), Formal analysis, Validation, Investigation, Resources, Visualization, Methodology, Writing—original draft, Writing—review and editing.

## Patent information

A patent application for the method described here has been filed.

## References

Agresti A. 2013. Categorical data analysis, 3rd edn. Wiley, Hoboken, NJ

Akaike H. 1974. A New Look at the Statistical Model Identification. IEEE Transactions on Automatic Control 19:716–723

Alpaydin E. 2014. Introduction to machine learning. The MIT Press, Cambridge, Massachusetts

Baraas RC, Zele AJ. 2016. Psychophysical Correlates of Retinal Processing. In: Kremers J, Baraas RC, Marshall NJ, eds. Human Color Vision. Springer International Publishing, Cham

Berg B, Cortazar B, Tseng D, Ozkan H, Feng S, Wei Q, Chan RY-L, Burbano J, Farooqui Q, Lewinski M, Di Carlo D, Garner OB, Ozcan A. 2015. Cellphone-Based Hand-Held Microplate Reader for Point-of-Care Testing of Enzyme-Linked Immunosorbent Assays. ACS nano 9:7857–7866. doi: 10.1021/acsnano.5b03203, PMID: 26159546

Chen Z, Liu J, Zeng M, Wang Z, Yu D, Yin C, Jin L, Yang S, Song B. 2012. Dot immunobinding assay method with chlorophyll removal for the detection of southern rice black-streaked dwarf virus. Molecules (Basel, Switzerland) 17:6886–6900. doi: 10.3390/molecules17066886, PMID: 22669043

Crowther JR. 2009. The ELISA guidebook, 2nd edn. Humana; [London : Springer, Totowa, N.J.

Davison AC, Hinkley D v. 1997. Bootstrap methods and their application. Cambridge University Press, Cambridge

Dorn PL, Engelke D, Rodas A, Rosales R, Melgar S, Brahney B, Flores J, Monroy C. 1999. Utility of the polymerase chain reaction in detection of Trypanosoma cruzi in Guatemalan Chagas’ disease vectors. The American journal of tropical medicine and hygiene 60:740–745. doi: 10.4269/ajtmh.1999.60.740, PMID: 10344645

Fisa R, Gállego M, Riera C, Aisa MJ, Valls D, Serra T, Colmenares M de, Castillejo S, Portús M. 1997. Serologic diagnosis of canine leishmaniasis by dot-ELISA. Journal of veterinary diagnostic investigation 9:50–55. doi: 10.1177/104063879700900109, PMID: 9087925

Henry Sum MS, Yee SF, Eng L, Poili E, Lamdin J. 2017. Development of an Indirect ELISA and Dot-Blot Assay for Serological Detection of Rice Tungro Disease. BioMed research international 2017:3608042. doi: 10.1155/2017/3608042, PMID: 29201901

Hosmer DW, Lemeshow S, Sturdivant RX. 2013. Applied logistic regression. Wiley, Hoboken New Jersey

Krudy Á, Ladunga K. 2001. Measuring Wavelength Discrimination Threshold Along The Entire Visible Spectrum. Periodica Polytechnica Ser. Mech. Eng. 45:41–48

Lacroix C, Renner K, Cole E, Seabloom EW, Borer ET, Malmstrom CM. 2016. Methodological Guidelines for Accurate Detection of Viruses in Wild Plant Species. Applied and environmental microbiology 82:1966–1975. doi: 10.1128/AEM.03538-15, PMID: 26773088

Lardeux F, Aliaga C, Depickère S. 2016. Bias due to methods of parasite detection when estimating prevalence of infection of Triatoma infestans by Trypanosoma cruzi. Journal of vector ecology : journal of the Society for Vector Ecology 41:285–291. doi: 10.1111/jvec.12224, PMID: 27860015

Lathwal S, Sikes HD. 2016. Assessment of colorimetric amplification methods in a paper-based immunoassay for diagnosis of malaria. Lab on a chip 16:1374–1382. doi: 10.1039/c6lc00058d, PMID: 27001468

Manoussopoulos IN, Tsagris M. 2015. Native Electrophoresis and Western Blot Analysis (NEWeB): Methods and Applications. Methods in molecular biology (Clifton, N.J.) 1312:343–353. doi: 10.1007/978-1-4939-2694-7_35, PMID: 26044016

McKinney SM, Sieniek M, Godbole V, Godwin J, Antropova N, Ashrafian H, Back T, Chesus M, Corrado GC, Darzi A, Etemadi M, Garcia-Vicente F, Gilbert FJ, Halling-Brown M, Hassabis D, Jansen S, Karthikesalingam A, Kelly CJ, King D, Ledsam JR, Melnick D, Mostofi H, Peng L, Reicher JJ, Romera-Paredes B, Sidebottom R, Suleyman M, Tse D, Young KC, Fauw J de, Shetty S. 2020. International evaluation of an AI system for breast cancer screening. Nature 577:89–94. doi: 10.1038/s41586-019-1799-6, PMID: 31894144

Nath S, Sarcar S, Chatterjee B, Chourashi R, Chatterjee NS. 2018. Smartphone camera-based analysis of ELISA using artificial neural network. IET Computer Vision 12:826–833. doi: 10.1049/iet-cvi.2017.0585

Oliphant TE. 2007. Python for Scientific Computing. Computing in science & engineering 9:10–20. doi: 10.1109/MCSE.2007.58

Olkkonen M, Ekroll V. 2016. Color Constancy and Contextual Effects on Color Appearance. In: Kremers J, Baraas RC, Marshall NJ, eds. Human Color Vision. Springer International Publishing, Cham

Pappas MG. 1994. The Biotech Business Handbook. Springer Science+Business Media, LLC, New York

R Core Team. 2018. R: A Language and Environment for Statistical Computing. R Foundation for Statistical Computing, Vienna, Austria

Racki N, Dreo T, Gutierrez-Aguirre I, Blejec A, Ravnikar M. 2014. Reverse transcriptase droplet digital PCR shows high resilience to PCR inhibitors from plant, soil and water samples. Plant methods 10:42. doi: 10.1186/s13007-014-0042-6, PMID: 25628753

Raschka S, Mirjalili V. 2017. Python machine learning: Machine learning and deep learning with Python, scikit-learn, and TensorFlow / Sebastian Raschka & Vahid Mirjalili. Packt Publishing, Birmingham, UK

Reinhard E. 2008. Color imaging: Fundamentals and applications.

A.K. Peters, Wellesley Mass Rodkvamtook W, Zhang Z, Chao C-C, Huber E, Bodhidatta D, Gaywee J, Grieco J, Sirisopana N, Kityapan M, Lewis M, Ching W-M. 2015. Dot-ELISA Rapid Test Using Recombinant 56-kDa Protein Antigens for Serodiagnosis of Scrub Typhus. The American journal of tropical medicine and hygiene 92:967–971. doi: 10.4269/ajtmh.14-0627, PMID: 25802430

Roggero P, Ciuffo M, Vaira AM, Accotto GP, Masenga V, Milne RG. 2000. An Ophiovirus isolated from lettuce with big-vein symptoms. Archives of virology 145:2629–2642, PMID: 11205109

Savary S, Ficke A, Aubertot J-N, Hollier C. 2012. Crop losses due to diseases and their implications for global food production losses and food security. Food Security 4:519–537. doi: 10.1007/s12571-012-0200-5

Schneider CA, Rasband WS, Eliceiri KW. 2012. NIH Image to ImageJ: 25 years of image analysis. Nature methods 9:671–675. doi: 10.1038/nmeth.2089

Schrader C, Schielke A, Ellerbroek L, Johne R. 2012. PCR inhibitors - occurrence, properties and removal. Journal of applied microbiology 113:1014–1026. doi: 10.1111/j.1365-2672.2012.05384.x, PMID: 22747964

Sing T, Sander O, Beerenwinkel N, Lengauer T. 2005. ROCR: visualizing classifier performance in R. Bioinformatics (Oxford, England) 21:3940–3941. doi: 10.1093/bioinformatics/bti623, PMID: 16096348

Smejkal GB, Kaul AC. 2001. Stability of Nitroblue Tetrazolium-based Alkaline Phosphatase Substrates. The Journal of Histochemistry & Cytochemistry 49:1189–1190

Subramanian S, Mahadevan A, Satishchandra P, Shankar SK. 2016. Development of a dot blot assay with antibodies to recombinant “core” 14-3-3 protein: Evaluation of its usefulness in diagnosis of Creutzfeldt-Jakob disease. Annals of Indian Academy of Neurology 19:205–210. doi: 10.4103/0972-2327.176867, PMID: 27293331

Suther C, Moore MD. 2019. Quantification and discovery of PCR inhibitors found in food matrices commonly associated with foodborne viruses. Food Science and Human Wellness 8:351–355. doi: 10.1016/j.fshw.2019.09.002

Trevethan R. 2017. Sensitivity, Specificity, and Predictive Values: Foundations, Pliabilities, and Pitfalls in Research and Practice. Frontiers in public health 5:307. doi: 10.3389/fpubh.2017.00307, PMID: 29209603

Tuszynski J. 2018. caTools: Tools: moving window statistics, GIF, Base64, ROC AUC, etc

van der Walt S, Colbert SC, Varoquaux G. 2011. The NumPy Array: A Structure for Efficient Numerical Computation. Computing in science & engineering 13:22–30.doi: 10.1109/MCSE.2011.37

Venkataramana M, Rashmi R, Uppalapati SR, Chandranayaka S, Balakrishna K, Radhika M, Gupta VK, Batra HV. 2015. Development of sandwich dot-ELISA for specific detection of Ochratoxin A and its application on to contaminated cereal grains originating from India. Frontiers in microbiology 6:511. doi: 10.3389/fmicb.2015.00511, PMID: 26074899

Waner T, Naveh A, Wudovsky I, Carmichael LE. 1996. Assessment of maternal antibody decay and response to canine parvovirus vaccination using a clinic-based enzyme-linked immunosorbent assay. Journal of veterinary diagnostic investigation 8:427–432. doi: 10.1177/104063879600800404, PMID: 8953526

Wes McKinney. 2010. Data Structures for Statistical Computing in Python. In: Stéfan van der Walt, Jarrod Millman, eds. Proceedings of the 9th Python in Science Conference

Wickham H. 2016. Ggplot2: Elegrant graphics for data analysis / Hadley Wickham; with contributions by Carson Sievert. Springer, Switzerland

World Health Organization. 2018. One Health. http://www.who.int/features/qa/one-health/en/

